# Occipital and parietal cortex participate in a cortical network for transsaccadic discrimination of object shape and orientation

**DOI:** 10.1101/2021.03.29.437597

**Authors:** B. R. Baltaretu, W. Dale Stevens, E. Freud, J. D. Crawford

## Abstract

Saccades change eye position and interrupt vision several times per second, necessitating neural mechanisms for continuous perception of object identity, orientation, and location. Neuroimaging studies suggest that occipital and parietal cortex play complementary roles for transsaccadic perception of intrinsic versus extrinsic spatial properties, e.g., dorsomedial occipital cortex (cuneus) is sensitive to changes in spatial frequency, whereas the supramarginal gyrus (SMG) is modulated by changes in object orientation. Based on this, we hypothesized that both structures would be recruited to simultaneously monitor identity *and* orientation across saccades. To test this, we merged two previous neuroimaging protocols: 21 participants viewed a 2D object and then, after sustained fixation or a saccade, judged whether the shape *or* orientation of the represented object changed. We, then, performed a bilateral region-of-interest analysis on identified cuneus and SMG sites. As hypothesized, cuneus showed both saccade and feature (i.e., orientation vs. shape change) modulations, and right SMG showed saccade-feature interactions. Further, the cuneus activity time course correlated with several other cortical saccade / visual areas, suggesting a ‘functional network’ for feature discrimination. These results confirm the involvement of occipital / parietal cortex in transsaccadic vision and support complementary roles in spatial versus identity updating.

## Introduction

The ability to extract pertinent visual information from our surroundings depends on the brain’s ability to aim, and account for, eye position [1–3]. Rapid eye movements (saccades) help gather new visual information by aligning the fovea with objects of interest, at the cost of disrupting visual stability and memory by displacing the retinal image relative to the world [4,5]. This necessitates some neural mechanism to continuously track both intrinsic object properties – cues to identity – and extrinsic spatial properties, like object location and orientation [5–8]. It is thought that during saccades, oculomotor signals are used to ‘update’ retinal location [9–15] and other visual features [5,7]. Recently, it has been shown that parietal and occipital cortex are involved in transsaccadic memory of spatial orientation and frequency, respectively [16–18]. However, in real world conditions, the brain must track intrinsic and extrinsic object properties simultaneously [19,20].

Various structures in the occipito-parieto-frontal stream have been implicated in the spatial aspects of transsaccadic vision, notably the parietal and frontal eye fields [9,10,21–26]. Likewise, transcranial magnetic stimulation (TMS) over various occipital, parietal, and frontal sites disrupts transsaccadic updating of object orientation [8,27–29]. More recently, we employed functional magnetic resonance imaging (fMRI) adaptation [30,31] to identify areas that are specifically involved in transsaccadic comparisons of object orientation. These experiments identified an area in the right supramarginal gyrus (SMG), anterolateral to the parietal eye fields, that showed transsaccadic orientation processing for both perception and grasp [16,17].

It has proven more difficult to localize the cortical mechanisms that update features related to object identity [32]. Modest amounts of feature remapping have been observed in monkey parietal eye fields [33]. A recent human study has shown that stimulus features (specifically, spatial frequency) can be decoded from whole-brain magnetoencephalography signals after an intervening saccade [34]. When we adapted our transsaccadic fMRI adaption paradigm to test spatial frequency, the main source of these modulations appeared to be superior-medial occipital cortex, i.e., cuneus [18]. Based on these results and our earlier experiments [16,17], we speculated that parietal and occipital cortex show a relative specialization for updating extrinsic spatial attributes of objects versus cues to intrinsic object identity, respectively.

In our previous adaption experiments [16–18], we only examined transsaccadic changes in one feature at a time. However, as noted above, in real world conditions, the brain must update multiple object features across saccades in order to track both object identity and its location / orientation. This, in turn, may require the ‘binding’ of multiple object features across saccades [35–37]. There is evidence that comparison between features evokes different mechanisms than the sum of the parts [38], but it is not known how multiple features are retained, compared, and discriminated across saccades.

Here, we investigated the cortical mechanisms for transsaccadic perception of multiple object features, combining intrinsic (identity-related) and extrinsic (spatial) features. To do this, we combined elements of our previous experiments [16–18]. Specifically, we used an fMRI paradigm that required participants to remember both object shape and orientation across a saccade and then discriminate which one of these had changed (Fig. 1). Our hypotheses were that 1) *both* SMG and cuneus would be modulated in this task, 2) SMG would show the saccade-orientation interactions that we observed previously [17] and 3) if cuneus is involved in object identity updating, our previous spatial frequency results should extend to the transsaccadic shape manipulation in the current task. To test this, we performed a bilateral region-of-interest (ROI) analysis, based on the sites obtained from our previous studies [17,18]. Our results confirmed the expected transsaccadic feature modulations in both SMG and cuneus, and suggest that the latter participates in a saccade-dependent network spanning several cortical lobes.

**Figure 1.**
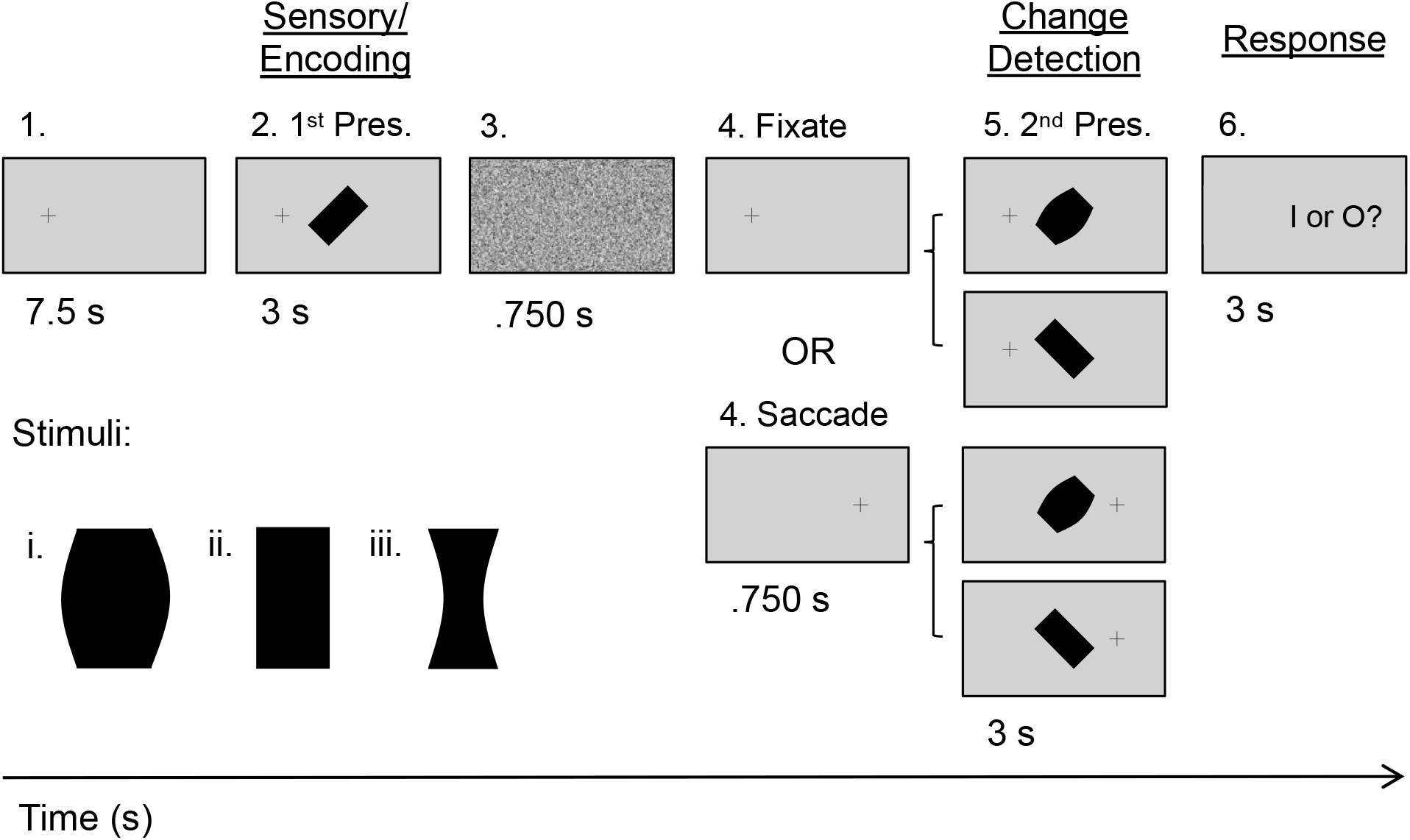
Experimental paradigm and criteria for transsaccadic feature modulation. An example trial is presented (initial left fixation, rectangle, 45°) with the four possible main conditions: 1) Fixation, Different Orientation, 2) Fixation, Different Shape, 3) Saccade, Different Orientation, or 4) Saccade, Different Shape. Each of the 19.5 s trials had three main phases: 1) *Sensory/Encoding*, during which the stimulus (one of the three possible objects: rectangle, barrel, or hourglass) was first presented at 45° to the left or right of vertical, while the participant was looking to the left or right of centre; 2) *Change Detection*, wherein the same object was presented at the orthogonal orientation (*Different Orientation* condition) or a different object at the same initial orientation (*Different Shape* condition) while participants fixated on the initial fixation cross (*Fixation* condition) or after they made a saccade (*Saccade* condition); and 3) *Response*, where participants made a button press response to make a judgment about whether the shape (‘I’) or the orientation (‘O’) of the object has changed across the two stimulus presentations.

## Results

To identify areas involved in transsaccadic discrimination of object orientation versus shape, we employed a task (Fig. 1) where 21 participants briefly viewed an object with one of three shapes (a rectangle, a convex ‘barrel’, or a concave ‘hourglass’) at one of two orientations (45° clockwise or counterclockwise). They either maintained fixation to the left or right of this central object or made a saccade to the opposite side. Then, they re-viewed an object with either a different shape or different orientation from among the same options, and then indicated which feature had changed. 17 participants met our behavioural inclusion criteria for analysis of fMRI data collected during this task.

Based on previous experiments [16–18], we expected both SMG and cuneus to be modulated in this task. To test this, we performed a hypothesis-driven region of interest (ROI) analysis on these sites (see next sections for details), and then performed a functional connectivity analysis to identify how saccade-related activity (if any) in these regions correlates with activity in other cortical areas.

### Region-of-interest analysis (I): Supramarginal Gyrus

In a previous experiment, we found that an area in right SMG, slightly anterior and lateral to the classic parietal eye field (revealed by a saccade-fixation subtraction) showed transsaccadic repetition modulations for object orientation, i.e., different activation when saccades were preceded and then followed by the same or different object orientations [16]. More recently, we showed that right SMG shows an interaction between this effect and saccades, i.e., significantly more orientation modulation during saccades compared to fixation [17]. Here, we predicted the same result in an ROI analysis based on the Talairach coordinates of peak right SMG activation in our previous study (Table 1) and the mirror left hemisphere site, forming a bilateral pair.

**Table 1.** Regions-of-interest, Talairach coordinates, t-statistic (T), and p-values (P) for *a priori* selected areas tested for saccade and feature modulations.

| Region | Talairach coordinates |  |  | Criterion 1 |  | Criterion 2 |  | Ref's |
| --- | --- | --- | --- | --- | --- | --- | --- | --- |
|  | x | y | z | T | P | T | P |  |
| <u>Occipital Cortex</u> |  |  |  |  |  |  |  |  |
| Cuneus | ±6 | -79 | 15 | 5.8 | 0.00028 | 2.9 | 0.0085 | [18] |
| <u>Parietal Cortex</u> |  |  |  |  |  |  |  |  |
| LH Supramargi-<br>nal Gyrus | -49 | -40 | 48 | -0.6 | 0.87 | 0.2 | 0.93 | [17] |
| RH Supramargi-<br>nal Gyrus | 49 | -40 | 48 | -0.2 | 0.93 | 0.1 | 0.93 | [17] |

Figure 2a, left panel, shows the location of the right SMG site relative to visual field-specific areas (Left Visual Field – Right Visual Field Stimulation across all (fixation and saccade) trials; yellow) and saccade-specific areas (Saccade – Fixation trials, all conditions; magenta), superimposed on an ‘inflated brain’. (Note that this rendering is useful for showing full brain data, but causes some spatial distortions and cluster separation.) Our right SMG site was anterior to two magenta clusters that might be part of the parietal eye fields, but did not itself show significant independent modulations for saccade or visual field. A whole-brain analysis of feature (i.e., shape) sensitivity at the initial fixation point (e.g., barrel versus hourglass, clockwise versus counterclockwise orientation) did not reveal significant clusters.

**Figure 2.**
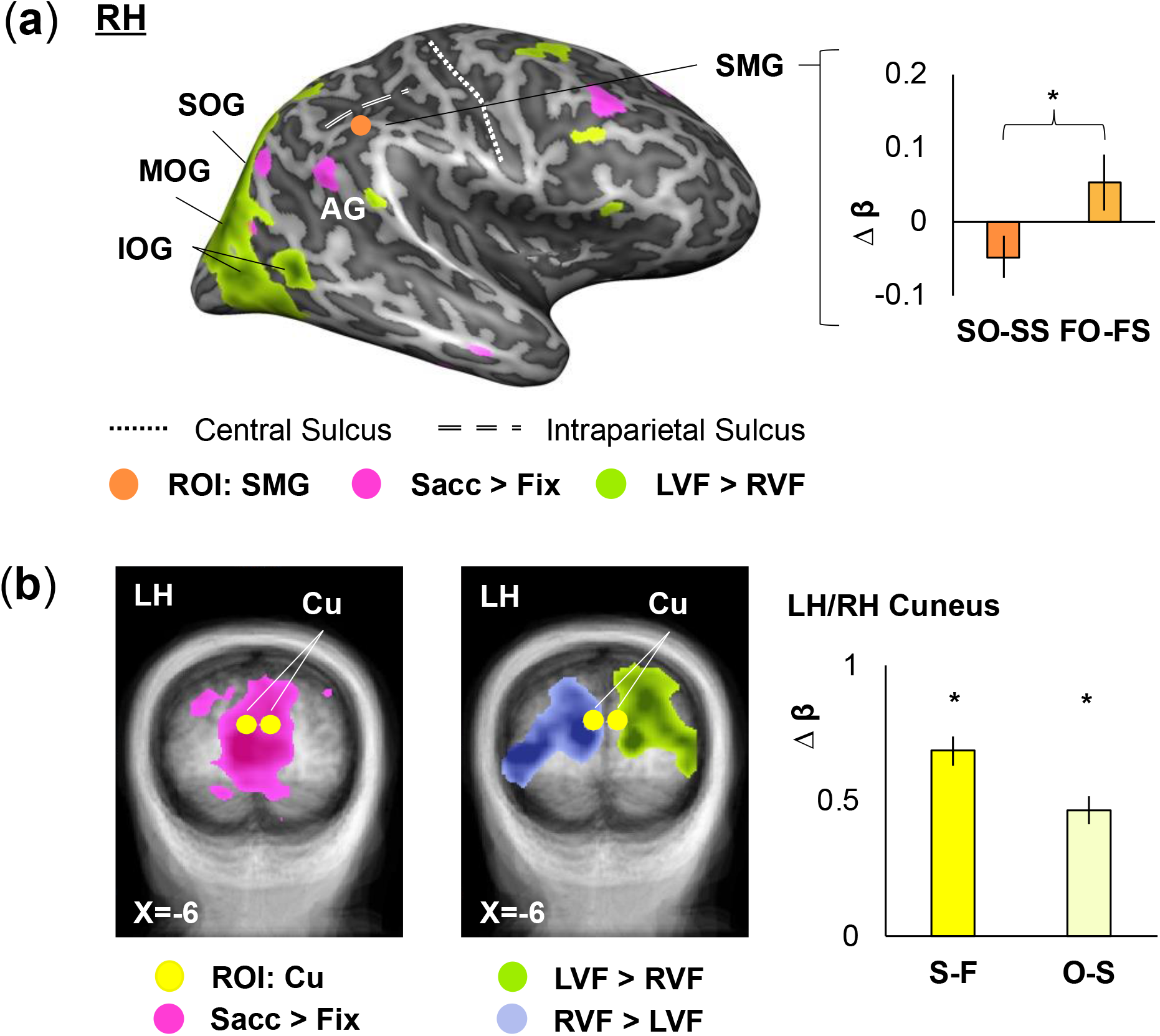
Region-of-interest (ROI) analysis results for tests of eye movement and feature effects. (**a**) Voxelwise maps (n=17) from two contrasts used to determine simple eye movement (Saccade > Fixation; “Sacc > Fix”; fuchsia) and visual field (Left Visual Field > Right Visual Field; “LVF > RVF” and “RVF > LVF”; lime green and cornflower blue) effects are overlaid onto inflated brain rendering of example participant’s right hemisphere (lateral view). Saccade sensitivity is shown parietally in angular gyrus (AG), whereas visual field effects can be observed in occipital cortex (extending from inferior through middle to superior occipital gyrus; IOG, MOG, and SOG). These regions are posterior/medial to right SMG (orange). B-weight differences depicted for right SMG in the bar graph on the right (error bars: SEMs) which show the significant interaction between eye movement- and feature-related changes across saccades (SO: Saccade, Orientation change condition; SS: Saccade, Shape change condition; FO: Fixation, Orientation change condition; FS: Fixation, Shape change condition). (**b**) Transverse slices (Talairach X-coordinates provided) for left and right cuneus (yellow) that show saccade (leftmost) and visual field (middle) sensitivity. B-weights differences (error bars: SEMs) from left (LH) and right (RH) cuneus are plotted (*left bar*: S: Saccade condition; F: Fixation condition; *right bar*: O: Orientation change condition; S: Shape change condition) and show significant eye movement and feature effects, as was expected from similar results in another transsaccadic perception study [18].

To test our specific hypothesis, we created spheres (radius = 5 mm) around our two SMG coordinates, and then extracted β-weights from these ROIs. We then performed the same interaction test as in our previous study on each ROI, i.e., Saccade (Same – different orientation) – Fixation (same – different orientation) [18], except in this case ‘same orientation’ corresponded to a shape change, thus: Saccade (Orientation change – Shape change) – Fixation (Orientation change – Shape change). SMG showed a significant hemispheric difference (F_1,16_ = 54.84, p = 1.5 × 10-6, η^2^ = 0.72), so we analyzed the left and right ROIs separately. RM-ANOVAs were then used to test our specific criteria in each of the ROIs. As predicted, right SMG passed this test (Fig. 2a, *right panel*) with a significant saccade – feature change interaction (F_1,16_ = 6.84, p = 0.019, η^2^ = 0.076). It is noteworthy, however, that the interaction was opposite to what we observed with orientation alone. Left SMG (not shown) did not show significant modulation for saccades (F_1,16_ = 0.39, p = 0.54, η^2^ = 0.010), feature (F_1,16_ = 0.048, p = 0.83, η^2^ = 8.5 η 10^−4^), or saccade – feature interactions (F_1,16_ = 1.46, p = 0.24, η^2^ = 0.025).

### Region-of-interest analysis (II): Cuneus

In a previous experiment [18], we found that right and left cuneus showed significant saccade modulations and right cuneus showed a significant transsaccadic feature modulation for *spatial frequency* (with left cuneus showing a similar trend). Neither area showed an interaction effect (perhaps because that occurs at some higher level). If this area plays a more general role in transsaccadic perception of object identity, we expected to see the same modulations for shape change in the current task.

To test this, we followed similar procedures and statistics as used for SMG. Figure 2b, left and middle panels, show locations of the cuneus ROIs relative to saccade and field-specific activity, respectively, superimposed on coronal slices. Both sites fall within a zone that contains both significant saccade and field-specific modulations. The right panel of Figure 2b shows the results of our specific hypotheses tests. In this case, cuneus did not show significant hemispheric differences (p > 0.05), so data from the bilateral pair were pooled. As predicted, this bilateral pair showed a significant main effect of eye movement (| S-F | F_1,16_ = 32.55, p = 3.25 × 10^−5^, η^2^ = 0.386) and feature (| O-S | F_1,16_ = 11.68, p = 0.015, η^2^ = 0.011). In this case, the directionality (analogous to same-different shape) was the same as what we observed for spatial frequency in a previous study [18]. Further, as in that previous study, we did not find a saccade-feature interaction (F_1,16_ = 0.54, p = 0.47, η^2^ = 0.003).

### Functional connectivity of cuneus with visual and sensorimotor regions

In the final step of our analysis, we performed a functional connectivity analysis based on our right cuneus ROI. To be consistent with our previous studies (and be feature agnostic), we based this analysis on the Saccade-Fixation subtraction [17,18]. This excluded SMG (since it did not show significant saccade modulations in this dataset). And, to be consistent with our previous study [18], we chose the right cuneus as the seed region. We then defined a sphere (radius = 5 mm) around this site and performed a psychophysiological interaction (PPI) analysis. This should identify a ‘network’ of cortical regions with similar saccade-related signals in this task.

This analysis revealed significant connectivity with (right) occipital, occipitotemporal, parietal, and frontal cortex (Fig. 3, Table 2). Specifically, we observed significant functional connectivity with early-to-intermediate visual occipital (right calcarine sulcus, CalcS, and superior occipital gyrus, SOG, as well as bilateral lingual gyrus, LG), sensorimotor parietal (right superior parietal lobule, SPL), oculomotor (left medial superior frontal gyrus, SFG, and frontal eye field, FEF) and spatial coding (right inferior frontal sulcus, IFS) frontal cortex.

**Table 2.** Functional connectivity network regions and Talairach coordinates resulting from psychophysiological interaction (PPI) analysis with right cuneus (seed region).

| Region | Talairach coordinates |  |  |  |  |  | Active Voxels | References |
| --- | --- | --- | --- | --- | --- | --- | --- | --- |
|  | x | y | z | Std x | Std y | Std z |  |  |
| <u>Seed Region</u> |  |  |  |  |  |  |  |  |
| RH Cuneus | 6 | -79 | 15 |  |  |  |  |  |
| <u>Occipital Areas</u> |  |  |  |  |  |  |  |  |
| LH Lingual Gyrus | -11 | -76 | -6 | 2.8 | 2.8 | 2.9 | 958 | [49] |
| LH Superior Occipital Gyrus | -28 | -77 | 14 | 2.6 | 2.5 | 2.1 | 587 | [50] |
| RH Calcarine Sulcus | 4 | -82 | -2 | 2.9 | 2.9 | 2.9 | 1000 | [51] |
| RH Lingual Gyrus | 3 | -86 | -14 | 2.9 | 2.8 | 2.8 | 980 | [49] |
| <u>Parietal Areas</u> |  |  |  |  |  |  |  |  |
| LH Superior Parietal Lobule | -19 | -77 | 39 | 2.8 | 1.6 | 2.5 | 437 | [52] |
| RH Superior Parietal Lobule | 9 | -68 | 55 | 2.7 | 2.1 | 1.8 | 379 | [53] |
| <u>Frontal Areas</u> |  |  |  |  |  |  |  |  |
| LH Frontal Eye Field | -25 | -12 | 53 | 1.3 | 1.2 | 2.2 | 141 | [54] |
| LH Superior Frontal Gyrus | -4 | 0 | 52 | 1.8 | 1.4 | 1.6 | 154 | [54] |
| RH Inferior Frontal Sulcus | 41 | 3 | 31 | 2.8 | 2.2 | 2.4 | 602 | [55] |

**Figure 3.**
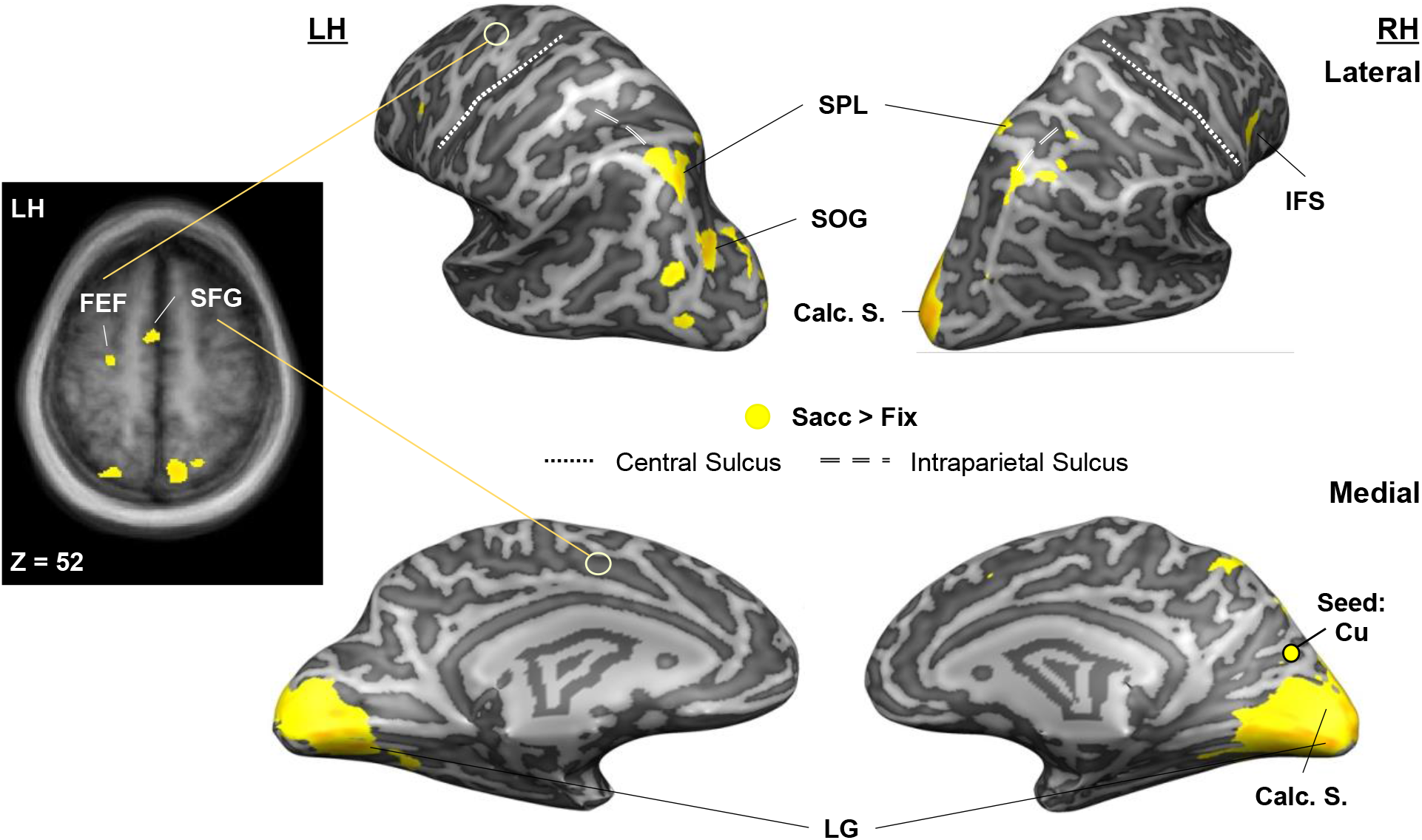
Functional connectivity network involved in transsaccadic updating of object features (orientation, shape). Using a Saccade > Fixation contrast, a psychophysiological interaction (PPI) contrast revealed statistically significant (RFX GLM; FDR (q < 0.05)) functional connectivity between right cuneus and frontal (right inferior frontal sulcus, IFS, left superior frontal gyrus, SFG, and frontal eye field, FEF), parietal (bilateral superior parietal lobule, SPL), and occipital (right calcarine sulcus, CalcS, left superior occipital gyrus, SOG, and bilateral lingual gyrus, LG) regions. (Note that these correlations do not necessarily imply direct anatomical connections.) Left: Transverse slice through the average brain of all participants (n=17) shows activation in left frontal (SFG, FEF) and right parietal (SPL) cortex in yellow which depicts correlation with the seed region, right Cu.

## Discussion

Our aim here was to investigate whether transsaccadic discrimination of multiple features relies on the same or different mechanisms as those for single features. Specifically, we tested if transsaccadic mechanisms related to intrinsic (spatial frequency) and extrinsic (orientation) object properties would combine and generalize to a task that involves discriminating object orientation versus shape changes. To do this, we tested for saccade and feature sensitivity in four regions that showed both these properties in previous transsaccadic tasks involving single features (SMG, cuneus) [17,18]. Of these regions, right SMG and bilateral cuneus showed the predicted patterns of saccade and feature sensitivity. Of these, cuneus showed main effects (of eye movement and feature change) and visual field sensitivity, whereas only SMG showed interactions, perhaps suggesting lower-level versus higher-level mechanisms. Further, saccade-related activity in cuneus correlated with activity throughout visual, control, and saccade-related regions of cerebral cortex. Overall, these analyses support the notion that SMG and cuneus participate in a complex network for transsaccadic retention of object identity and orientation.

### Monolithic versus Feature-specific transsaccadic mechanisms

Although some theoretical discussions of transsaccadic feature perception have suggested a role for occipital cortex [6,7,39,40], initial experimental studies emphasized the parietal cortex as the key hub [8,16,17,33]. However, considering the diversity and regionalization of visual functions [30,41,42], and the prevalence of saccades in normal vision [43,44], the notion that the combination of saccades and vision would be handled by one specialized mechanism seems naïve. As noted in the introduction, transsaccadic updating of object location is found throughout visual and visuomotor cortex [5,9,34,45], whereas transsaccadic perception of object orientation, spatial frequency, and shape appears to involve parietal, medial occipital, and in some cases, lateral occipital cortex [16–18]. The current results are also consistent with a more distributed system, with feature-specific modules.

### Role of SMG in transsaccadic perception

Although the orientation of local features (such as the lines that make up curves and shapes) can be a cue to identity [46–48], in our experiments, all lines in the stimuli have the same orientation (Fig. 1), and the task instructions strongly reinforced the interpretation that this was a cue to the orientation of the entire object. Given the consistency of our results across various experiments [6,16,17] (and the current experiment), it appears that right SMG plays a specific role in updating this property across saccades. (See those previous papers and ‘networks’ below for discussions of the hemispheric specialization of this structure). Taken together with the many studies of spatial updating (e.g., [10,22]), this suggests a general role for parietal cortex in the transsaccadic spatial updating of the fundamental extrinsic spatial properties of objects: location, orientation, and the combination of these two properties [6,7].

### Possible role of medial occipital cortex in transsaccadic perception

In this and our previous study on spatial frequency [18], dorsomedial occipital cortex appears to play a role in the postsaccadic integration of spatial frequency and object shape. The cuneus sites described here likely fall within a region spanning visual areas V2 and V3 [56]. These areas are ideally suited to provide the rudimentary machinery for transsaccadic feature analysis [6,7,57,58].

First, they are well-positioned to provide rapid, bottom-up operations on their anatomic inputs from V1 [59–61]. Second, they include the requisite machinery for feature orientation, spatial frequency, and early shape processing [62–64]. Third, there are indications that even these areas are able to store information in some sense [58]. Fourth, these areas – V1, V2, and V3 – are known to show saccade modulations consistent with remapping signals [23,29,57,58]. Finally, these areas have reciprocal connections with both ‘ventral stream’ object recognition areas [65–67], and dorsal stream saccade and attention areas [68–70]. This feedback, in combination with V1/V2/V3 retinotopic topography (absent at higher levels), makes this area also ideally suited for integrating location and identity information across saccades [6,7,39].

Thus, situated at both early and intermediate stages of object feature processing, with the capability to monitor several object feature changes and receive feedback signals for attention and saccades, medial occipital cortex is an excellent, additional candidate for transsaccadic feature perception. Our experiments suggest that this area might play a specific role in transsaccadic processing of lower-level intrinsic object features related to identity, presumably working in concert with higher level ‘ventral stream’ mechanisms like lateral occipital cortex and inferior parietal cortex [20,71,72].

### Combining mechanisms for extrinsic and intrinsic feature updating

As noted above, transsaccadic perception would be somewhat useless if we could not update and link both object identity and location / orientation across saccades [50,73]. There is evidence that comparison between features evokes different mechanisms than the sum of the parts [38]. Based on the common properties of spatial frequency and shape (as intrinsic object features related to object *identity)*, we speculated that both cuneus and SMG would be modulated by transsaccadic shape changes. In the experiment, both cuneus and SMG showed the expected modulations, although the saccade-feature interaction had the opposite directionality [17]. This suggests that the individual mechanisms for transsaccadic orientation and shape updating combined, although not necessarily in a linear fashion. What remains to be understood, is how these various signals are combined – the classic ‘binding problem’ [74,75]. While cuneus and SMG were not linked in the same functional network in the current study (likely because SMG did not show strong enough saccade modulations), there is enough overlap between their functional networks in different studies (see next section) to suggest that this is a possible linking mechanism.

### Transsaccadic networks: Task-specificity

Considering the complexity of transsaccadic signals discussed above, it seems likely that these mechanisms can be better understood at a network level. In our previous study, where participants were required to update grasp orientation across saccades [17], they recruited a cortical network that included each element required for the task, including a transsaccadic orientation updater (SMG), saccade signals (FEF), and grasp motor areas (anterior intraparietal and superior parietal cortex). Likewise, in the current study, we identified a network that seems eminently suitable for our feature discrimination task (Fig. 3): calcarine sulcus for early feature processing [56,64,76], lingual gyrus and superior occipital gyrus for mid-level shape recognition [77,78], posterior parietal cortex for saccade/reach updating signals [9,10,22,79], and oculomotor superior frontal gyrus (likely pre-supplementary motor area)/frontal eye field [27] and spatial coding in inferior frontal sulcus [80]. The differences in these studies might thus simply be due to task and feature details.

Finally, although we used right cuneus as a seed region here, it is noteworthy that its activity predominantly correlated with activity in right occipital and parietal cortex, with oculomotor activation in left hemisphere. Right cortex also predominated in our previous regional and network analyses [16–18]. Consistent with this, transsaccadic memory is most sensitive to right parietal stimulation and brain damage [6,52,81]. This is generally consistent with the role of parietal cortex in spatial attention and cognition and suggests that impaired transsaccadic perception might contribute to better-known disorders like hemispatial neglect [82].

## Conclusion

Here, we used fMRI to determine the cortical correlates of transsaccadic perception in a task where participants had to remember and discriminate changes in object orientation versus shape after a saccade. Our current findings support specific roles of two regions in transsaccadic feature discrimination: SMG for orientation and medial occipital cortex for shape. Taken together with the previous literature, these results suggest that the cortical mechanisms for transsaccadic perception are both feature- and task-dependent. Specifically, there seems to be relative specialization for updating of object location and orientation in parietal cortex [9,16–18,57,58] versus transsaccadic perception of intrinsic object features in medial occipital cortex (here and [18]). These areas appear to work in conjunction for more complex tasks, apparently forming ad-hoc functional networks with saccade centers (providing the updating signal) and higher order recognition and/or motor areas, as required for the task.

## Materials and Methods

### Participants

To determine the minimum number of participants required to attain adequate statistical power, we used the effect size from right SMG, which is closest to predicted SMG here, found in the most recent of a series of transsaccadic perception studies [17]. We performed a power analysis (G*Power) using this effect size (0.887), with the following parameters: 1) two-tailed t-tests for the difference of matched pair means, 2) a desired power value of 0.90, and 3) an alpha-value of 0.05. On this basis, the suggested number of participants was 13, though we wanted to ensure full desired power, so we recruited 21 participants for this study, of whom we analyzed data from 17 participants (see below). The expected power was 0.914.

Twenty-one participants with normal or corrected-to-normal vision from York University, Toronto, Canada took part in this study. One participant’s data were excluded, as this person did not complete the experiment, and three participants’ data were excluded due to poor behavioural performance (i.e., less than 80% behavioural accuracy across all runs), which left 17 participants’ data for all further analyses. Participants were compensated monetarily for their time. All participants were right-handed (mean age: 28.7 +/− 5.2, 12 females). All participants provided written informed consent and did not have any neurological disorders. This experiment was approved by the York University Human Participants Review Subcommittee.

### Experimental set-up and stimuli

Participants were first required to pass MRI screening. Upon successful completion of this step, participants were introduced to the task. Once they felt comfortable with the requirements of the task, they were asked to assume a supine position on the MRI table. Their head lay in a 64-channel head coil. A mount on the head coil was used to support an infrared eye-tracker (for the right eye) to ensure appropriate fixation/eye movements, as well as a mirror that reflected the image of the screen from inside the MRI bore. iViewX software (SensoMotoric Instruments) was used to record eye movements for offline verification of correct behavior (see exclusion criteria above). Participants were provided with a button box that was held with the right hand (index and middle fingers positioned over the first and second buttons, respectively). Button press responses were analyzed offline.

The experiment was conducted under conditions of complete darkness. A fixation cross was always visible (except during the presentation of the mask) to direct participants’ gaze as required by the task. The fixation cross could appear approximately 7° from either the left or right edge of the gray screen (45.1° × 25.1°) along the horizontal meridian. The black stimulus always appeared in the center of the screen and was either a: 1) rectangle, barrel-shaped object, or 3) hourglass-shaped object (Fig. 1a). The dimensions of the rectangle were 12° × 6° the other two objects had the same area as the rectangle.

### General paradigm/procedure

#### Experiment

To determine the cortical correlates of transsaccadic perception of object orientation versus shape, we used an event-related fMRI design (Fig. 1). Each trial commenced with a fixation cross, presented either to the left or right center, for 7.5 s. Then, the central object (rectangle, barrel-shaped, or hourglass-shaped object) appeared for 3 s at +/− 45° from vertical. This was followed by a static noise mask for 0.75 s to avoid an afterimage. The fixation cross would then appear either at the same position as before the mask (*Fixation* condition) or at the other location (*Saccade* condition) for 0.75 s. The same object presented in a different orientation (*Orientation change* condition) or one of the other two different objects presented in the same orientation (*Shape change* condition) appeared in the center for 3 s. The object disappeared and the fixation cross was replaced by an instruction (“I or O?”) for 3 s, indicating that participants should indicate if the two objects presented in the trial changed in shape (first button, ‘I’) or in orientation (second button, ‘O’). Thus, there were four main conditions: 1) Fixation, Orientation change, 2) Fixation, Shape change, 3) Saccade, Orientation change, or 4) Saccade, Shape change. These trial types were randomly intermingled within a run (24 trials); there were eight runs in total. Each run began and ended with central fixation for 18 s to establish baseline measures.

#### Imaging parameters

We used a 3T Siemens Magnetom Prisma Fit magnetic resonance imaging (MRI) scanner. Functional experimental data were acquired with an echo-planar imaging (EPI) sequence (repetition time [TR]= 1500 ms; echo time [TE]= 30 ms; flip angle [FA]= 78 degrees; field of view [FOV]= 220 mm × 220 mm, matrix size= 110 × 110 with an in-slice resolution of 2 mm × 2 mm; slice thickness= 2 mm, no gap). A total of 312 volumes of functional data (72 slices) were acquired in an ascending, interleaved manner for each of the eight runs. In addition to the functional scans, a T1-weighted anatomical reference volume was obtained using an MPRAGE sequence (TR= 2300 ms, TE= 2.26 ms; FA= 8 degrees; FOV= 256 mm × 256 mm; matrix size= 256 × 256; voxel size= 1 × 1 × 1 mm^3^). 192 slices were acquired per volume of anatomical data.

### Analysis

#### Behavioural data

Eye movements and button presses were recorded throughout the experiment; the former were monitored during the experiment and analyzed offline to ensure appropriate fixation and saccade production. If participants made eye movements when not required to, the data associated with those trials were excluded from further analysis. In addition, data for trials where incorrect button presses were made were excluded from any additional analysis. On these bases, 148 trials were removed (out of 3264 trials, representing 4.5%; maximum four trials were removed across all participants, with a mode of four) for the 17 participants.

#### Functional imaging data

Functional data from each run for each participant were preprocessed (slice time correction: cubic spline, temporal filtering: <2 cycles/run, and 3D motion correction: trilinear/sinc). Raw anatomical data were transformed to a Talairach template [83]. Functional data were co-registered via gradient-based affine alignment (translation, rotation, scale affine transformation). Finally, data were smoothed using a Gaussian kernel of 8 mm full width at half maximum. On the basis of missing predictors (more than 50% of trials with incorrect behavioral response), nine runs across all participants were removed from the overall random effects general linear model (RFX GLM; i.e., 216 trials out of 3116 trials; 6.9%).

Four separate GLMs were generated for each participant for every run. The first set of GLMs contained 4 predictors: 1) Fixation, Orientation change, for an entire trial during which participants maintained fixation throughout while only the orientation of the same object changed; 2) Fixation, Shape change, for a trial during which participants maintained fixation while only the shape of the object changed; 3) Saccade, Orientation change, for a trial during which participants had to make a saccade while only the orientation of the object changed; and 4) Saccade, Shape change, for a trial where participants made a saccade while only the shape of the object changed. The first presentation of the stimulus was referred to as the *Sensory/Encoding* phase and the second presentation period was referred to as the *Change Detection* phase. These GLMs were created in BrainVoyager QX 20.6 (Brain Innovation), where each of the predictors was convolved with a haemodynamic response function (standard two-gamma function model) [84]. GLMs were additionally adjusted via the addition of a last, confound predictor (“Error”) to account for any errors made during trials due to unwanted eye movements or incorrect button responses.

The second set of GLMs were structured such that the *Sensory/Encoding* phase was modeled with two predictors to ultimately determine visual field effects: 1) “LVF” for when the object appeared in the left visual field relative to fixation and 2) “RVF” for trials where the object appeared in the right visual field relative to fixation. The rest of the trial was modeled with “Fix” or “Sacc” predictors to indicate if it was a Fixate or Saccade trial, respectively. An “Error” confound predictor was included to model trials where eye movement or button press errors had been made.

The third set of GLMs were modified to test for the presence of orientation-sensitive bias during the first stimulus presentation. As such, two predictors were modeled (as before) for the *Sensory/Encoding* phase: 1) “45°” for objects that were initially presented at 45° and 2) “135°” for objects that were initially presented at 135° from horizontal. The remainder of the trial was simply coded according to whether it was a Fixate or Saccade trial with “Fix” or “Sacc” predictors. A confound predictor (“Error”) was also included for eye movement or button press errors.

The last set of GLMs were modified to determine whether there was an initial shape-related bias upon first stimulus presentation. Three predictors, one for each of the three object shapes, were used to model the *Sensory/Encoding* phase: 1) “Rect” for when the rectangle was presented during this phase; 2) “Barrel” for the barrel-shaped object; and “Hourglass” for the hourglass-shaped object. Similar to the two previous GLMs, the remainder of the trial was modeled according to whether it was a Fixate or Saccade trial (i.e., “Fix” or “Sacc” predictors, respectively). Finally, the “Error” predictor was also included for erroneous eye movements and/or button presses.

#### Region-of-interest (ROI) analysis

A hypothesis-driven region-of-interest (ROI) analysis was performed based on Talairach coordinates obtained our previous studies [17,18]. Specifically, we used the previously identified coordinates of peak modulation for transsaccadic orientation changes (right SMG) and spatial frequency changes (right cuneus), which we hypothesized would generalize to transsaccadic shape detection if cuneus plays a more general role in object *identity*. We then mirrored these coordinates laterally to obtain four ROIs (i.e., bilateral SMG and cuneus; Table 1). ROIs were created around these coordinates as spheres with 5-mm radius using BrainVoyager QX v2.8 (Brain Innovation). β-weights were then extracted and inspected for hemispheric differences with a two-tailed repeated-measures t-test. If a significant result was obtained, the β-weights from the left and right hemispheres were analyzed separately for each of the two criteria (see Results for details). On this basis, we pooled data only for cuneus. Within each of the ROIs, we ran a repeated-measures analysis-of-variance (RM-ANOVA) with two factors: Eye movement (2 levels: Fixation, Saccade) and Object feature (3 levels: Rectangle, Barrel, Hourglass). See Results and Table 1 for further information.

#### Whole-brain univariate analysis for saccade and visual field sensitivity

We performed a whole-brain analysis to determine the presence of saccade sensitivity within the ROIs. For these univariate analyses, we applied four independent contrasts to test for four effects. First, we performed a whole-brain voxelwise Saccade > Fixation contrast to identify cortical regions that were modulated by saccades in this task (Fig. 2). Second, we applied a Left > Right Visual Field contrast using the predictors for the *Sensory/Encoding* phase to find regions that are sensitive to perception of objects within the respective visual fields (Fig. 2). To each of the resulting volumetric maps, we applied False Discovery Rate correction to account for multiple comparisons (q < 0.05) BrainVoyager QX v2.8, Brain Innovation).

#### Psychophysiological interaction

Our last goal was to identify the functional network for saccade-related updating during transsaccadic perception. In order to do this, we conducted psychophysiological interaction (PPI) analysis [17,84–86] on data extracted from the cuneus peak site (Talairach coordinates: 6, −79, 15). For optimal signal processing, we used a sphere of radius = 5 mm around this site. To perform this analysis, we used three predictors: 1) a physiological component (which is accounted for by z-normalized time-courses obtained from the seed region for each participant for each run); 2) a psychological component (task predictors, i.e. for the Saccade and Fixation conditions, were convolved with a hemodynamic response function), and 3) a psychophysiological interaction component (multiplication of seed region z-normalized time-courses with task model in a volume-by-volume manner). For the psychological component, the *Saccade* predictors were set to a value of ‘+1’, whereas the Fixation predictors were set to a value of ‘-1’; all other predictors were set to a value of ‘0’ in order to determine the responses to Saccades as compared to Fixation. (For greater detail, please see O’Reilly et al. [86] and [17]). We created single design matrices for each run for each of the 17 participants, which were ultimately included in an RFX GLM. Then, we used the third, psychophysiological interaction predictor to determine the regions that constitute the functional network that shows saccade-related modulations for feature updating. To the resulting volumetric map, we applied FDR correction (q < 0.05).

## Contributions

We thank Saihong Sun for technical support and Joy Williams and Dr. Diana Gorbet for help with data collection.

